# Microtranscriptome of contrasting sugarcane cultivars in response to aluminum stress

**DOI:** 10.1101/645267

**Authors:** Renan Gonçalves Silva, Thiago Mateus-Rosa, Suzelei de Castro França, Pratibha Kottapalli, Kameswara Rao Kottapalli, Sonia Marli Zingaretti

## Abstract

Although metallic elements are required for plant growth, aluminum ions (Al^+3^) can be considered one of the major abiotic factors affecting productivity. In plants, the presence of Al^+3^ can result in inhibition of root growth triggering water and nutrient deficiency. Plants under stress conditions undergo gene expression changes in specific genes or post-transcriptional gene regulators as miRNAs that can led to resistance. In this study, we investigated the miRNAs involved in the sugarcane response to aluminum stress. Four miRNA libraries were generated using sugarcane roots of two contrasting (tolerant and sensitive) sugarcane cultivars growing under aluminum stress to identify the miRNAs involved in the sugarcane response. Here we present the first miRNAs sequencing of sugarcane response under aluminum stress. The contrast of the cultivars seen in the field was reflected in the micro transcriptome with opposing expression profile. We selected 394 differentially expressed miRNAs, in both cultivars, 22% were common between cultivars. Real time quantitative polymerase chain reaction was used to validate the differentially expressed miRNAs through high-throughput sequencing in sugarcane roots. Target genes prediction was also analyzed. Our results indicated miRNAs that modulated specific target genes involved in roots development and plant aluminum stress response. Those genes can be the answer to tolerance in sugarcane and used in breeding programs to develop tolerant cultivars.

## Introduction

Sugarcane (*Saccharum* spp.) as an important source of sugar and ethanol became the third most produced commodities in the world (1.4G). In this context, Brazil figure as a major sugarcane producer (500M tons) followed by India (300M), China and Thailand [1]. The projections, based on the worldwide increasing demand for food and energy, are that sugarcane global production will increase by 21% until 2024. Production can be increased by increasing productivity and cropland expansion. The sugarcane crop expansion is evident in Brazil, where nowadays more than 9.5 million ha are used to cultivate sugarcane, but the demand for sugar and ethanol will increase to 10.5 million ha by the years 2023/24 [2].

Among the main factors that can affect agricultural productivity, soil has fundamental importance since it offers not only physical support but also water and the necessary nutrients for plant growth. Aluminum (Al) together with silicon and oxygen are the three most abundant elements in earth crust. Although metallic elements are required for plant growth, aluminum ions (Al^+3^) can be considered one of the major abiotic factors affecting agriculture productivity [3]. Al is a nonessential element naturally found in the soil but it is toxic and its bioavailability is highest on acidic soils (pH of 5.5 or lower), resulting in inhibition of root growth, architecture alteration and elongation disruption [3]. Around the world 50% of arable soils are acidic [4], in Brazil acidic soil comprises 500 million hectares, and 70% of this land been used for sugarcane plantation [5].

Most of the Al^+3^ is accumulated in the root apoplast and then translocated to other tissues [6], and the action of Al^+3^ on roots and plant development depends on the exposure time and aluminum concentration. The effects of Al^+3^ on plant metabolic process can be seem just few minutes after plant been exposed. In plants exposed to Al^3+^ (1.4µM) it was detected in the nuclei inhibiting cell division and cell viability after 30 min (Silva et al., 2000). Due to the rapid reactivity of Al^+3^ the first changes occur in the cell wall, plasma membrane, cytoskeleton and the cell nucleus [7]. This process inhibits root growth and they become shorter and thicker, absorbing less nutrients and water, and transporting molecules more slowly through the cells [8, 9], triggering water stress and nutrient and mineral deficiency [10]. In sugarcane, the inhibition of roots growth can reach 46% under Al stress [11].

Plants under stress conditions can undergo gene expression changes that can led to resistance. Those changes can be specific functional gene expression or post-transcriptional gene regulation. It can be achieved by the expression of transcriptions factors (TFs) like MYB proteins, a key player in the regulation of plant response abiotic stress [12], or small noncoding RNAs named microRNAs (miRNAs) important gene regulators at post-transcriptional levels. miRNAs are single strand RNA sequence, 20 to 24 nucleotides long, and in plants they act in the pos-transcriptional gene silence (PTGS) level [13, 14, 15]. The first identified miRNAs were involved in modulating physiological and biochemical process that regulate plant development and adaptation [16]. The miRNAs identified in different plants such as: *Arabidopsis thaliana* [17], *Triticum aestivum* L. [18], *Glycine max* [19], *Manihot esculenta* [20, 21], suggest that miRNA also plays an important role in the regulation of molecular responses to biotic and abiotic stress.

Over the last years, miRNAs have been intensively studied but not much is known about metal stress plant response, especially in crop plants. The available information about aluminum stress plant responses comes from model plants such as *Medicago truncatula* [22, 23] and *Arabidopsis thaliana* [24, 25]. Under metal stress, the plant gene expression can be modified to regulate different mechanisms such complexation of excess metal, defense against oxidative stress and signal transduction for different biological process [26, 27]. Some miRNAs such as miR159, miR160, miR319, and miR396 had been identified as down-regulated in *Medicago truncatula* seedling roots after 4 hours of under aluminum stress, their targets are transcription factors related to seed germination, embryo development, cold and drought response [23].

In sugarcane several miRNAs associated to cold [28] and drought [29, 30, 31] tolerance were identified, however, there is no data of miRNAs involvement in response to Al stress. Our goal is to understand the molecular mechanisms of abiotic stress tolerance in sugarcane and the role of miRNA’s in this response to aluminum stress. In this study, we focused in differential miRNA expression analysis and quantitative real-time PCR (qRT-PCR) validation in sugarcane roots growing under increased level of aluminum (Al^3+^) to understand the molecular mechanisms of aluminum stress tolerance.

## Materials and methods

### Plant materials and RNA isolation

Pre-germinated plants from two sugarcane (*Saccharum* spp) cultivars, CTC-2, tolerant to aluminum stress (TAS) and RB-855453, sensitive to aluminum stress (SAS), were grown in a hydroponic system in a greenhouse at 26°C to 30°C range and natural dark/light cycles. For 30 days plants were kept in 16L container filled with standard hydroponic solution [32] before going under stress when plants were cultivated for seven days under two aluminum concentration (0.0 and 22Lµmol Al^+3^ L^−1^) and pH 4.5. After seven days, roots were collected and immediately frozen in liquid nitrogen and stored at −80°C for further use. Total RNA was isolated from control and stressed plants root samples using the Sigma plant RNA kit (Sigma, Inc, USA). RNA quality and concentration were determined by Qubit 2.0 fluorometer (Life Technologies, USA).

### miRNA library and sequencing

cDNA libraries were generated using Illumina True-Seq small RNA prep (Illumina, USA) and sequenced using 35bp single end sequencing on MiSeq sequencer (Illumina, Inc, USA) following the manufacture instruction.

### Real time PCR of miRNAs

In order to validate our miRNA transcriptome we performed a qPCR analysis of randomly selected miRNA using the Stem-loop and quantitative real time polymerase chain reaction (qRT-PCR) [33]. For cDNA synthesis, RevertAid First Strand cDNA Synthesis kit (Thermo Fisher Scientific, USA) was used following the manufacture instructions. For qRT-PCR experiments, cDNA concentration was standardized for each sample and dissociation curve analysis was performed to check primer specificity. The reaction was performed in 20 μL containing 1 μL of RNA, DNAse treated, 200U of RevertAid M-MuLV Reverse Transcriptase, 20 mM DNTPs, RiboLock RNase Inhibitor (20 U), 5X reaction buffer (Thermo Fisher Scientific, USA), RT specific Primer 1 μM, dT primer (100 μM), at 42°C for 60 minutes and 5 minutes at 70°C. Real time PCR was carried out in a Stratagene MX3005P thermocycler using SYBR Green Jump Start Taq Ready Mix (Sigma Aldrich, USA) for quantifying amplification results. Thermal cycling conditions were as follow: 94°C for 2 minutes followed by 40 cycles of 94°C for 15 s, 60°C for 1 minute and 72°C for 30 seconds.

The miRNAs expression levels were quantified after normalization to 18SrRNA gene used as internal control. The gene specific primers used in the real time experiments and miRNAs sequences are in S2 and S3 Tables. For the RT qPCR experiment two time points were used, initial and 7 days after stress (DAS). miRNAs expression levels were analyzed using MXPro qPCR software 4.10 version (Stratagene, USA). Three biological replicates were examined to ensure reproducibility.

### miRNA targets prediction and functional annotations

The targets of the miRNAs were predicted using Mercator (http://mapman.gabipd.org/web/guest/app/Mercator) by searching for targets genes based in the MapMan “BIN” ontology, which is tailored for functional annotation of plant "omics" data [34]. The GO (Gene Ontology) categorization were listed as three independent hierarchies for biological process, cellular component and molecular function using UniProt Knowledgebase (https://www.uniprot.org) and QuickGO (EMBL-EBI, https://www.ebi.ac.uk/QuickGO) tools. The data of individual biological library were deposited to NCBI SRA database with SRA accession IDs: SRR9035251, SRR9035250, SRR9035245, SRR9035244, SRR9035249, SRR9035248, SRR9035243, SRR9035242, SRR9035253, SRR9035252, SRR9035247 and SRR9035246.

## Results

### Construction and sequencing analysis of miRNAs library

To identify the miRNAs involved in the aluminum stress response four-miRNA libraries, generated from the sugarcane roots of two contrasting sugarcane cultivars CTC-2 (Tolerant Aluminum Stress, TAS) and RB-855453 (Sensitive Aluminum Stress, SAS), under aluminum stress for seven days, were sequenced using Illumina technology. Over 12 million raw reads, with a Q-Score of 37 and 53% CG content, was obtained. After processing and filtering for poor quality sequence, 5.8 million from the CTC-2 (TAS) and 6.2 million reads from RB-855453 (SAS), clean sequences remained. About 20K reads were assembled, 11.5K from RB-855453 (SAS) and 8.5K from CTC-2 (TAS). The size distribution of the miRNAs ranged from 17 to 28 nt, as it is been presented in (Fig 1). The majority of the reads were from 20 to 24nt in length with 21nt being the most redundant species for both cultivars. The size distribution of sugarcane roots small RNAs is consistent with results observed in other plants using a deep-sequencing approach [35, 36].

**Fig 1.**
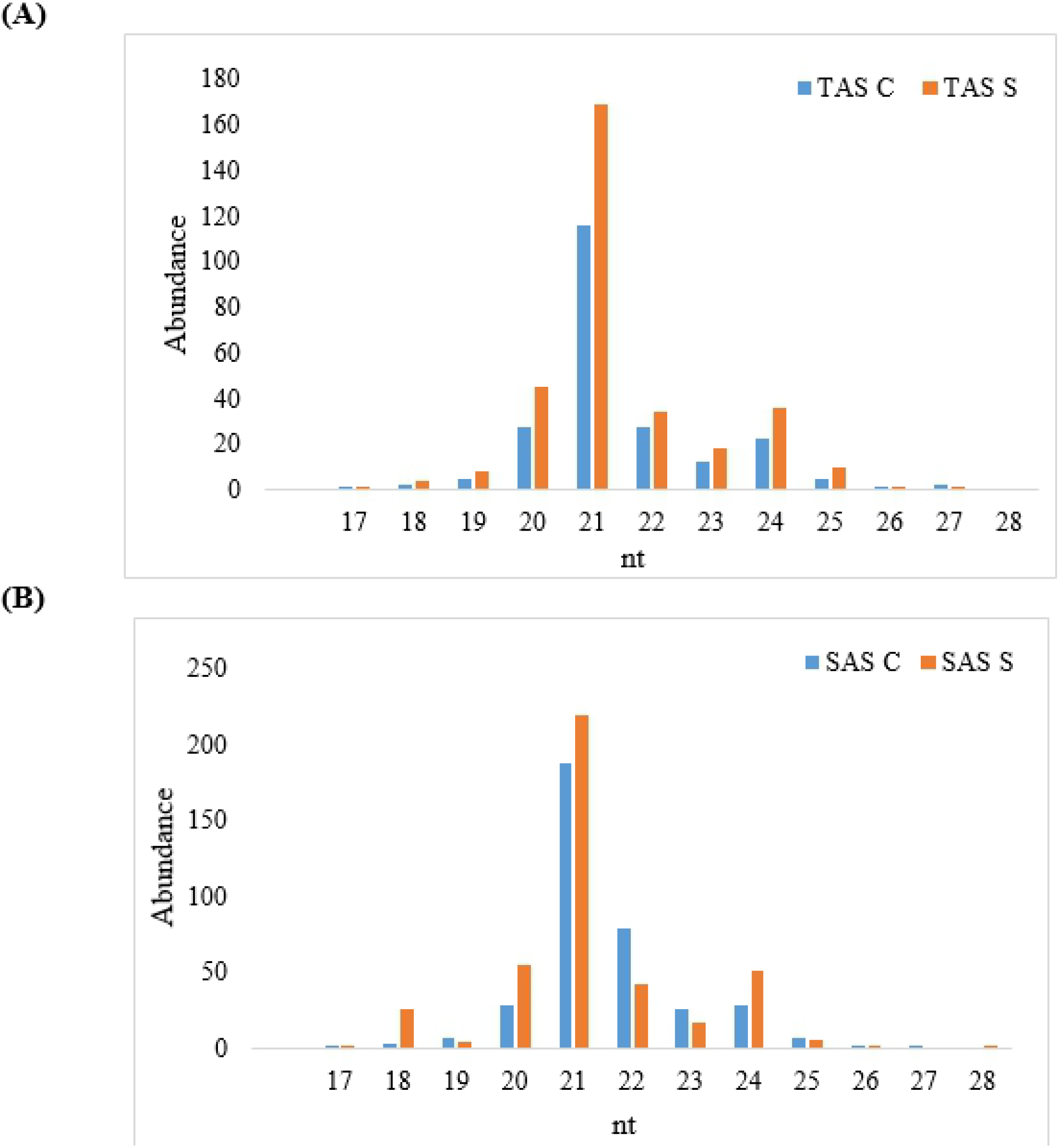
Size distribution of miRNAs sequences in two sugarcane cultivars. (A) Abundance in tolerant cultivar. TAS C – Tolerant aluminum stress control; TAS S– Tolerant aluminum stress stressed; (B) Abundance in sensitive cultivar. SAS C – Sensitive aluminum stress control; SAS S – Sensitive aluminum stress stressed.

To identify the miRNAs involved in the sugarcane response to aluminum stress we selected the miRNAs differently expressed in both cultivars. A total of 394 differentially expressed miRNAs were identified (S1 Table); 104 were specifically in TAS and 116 specifically in SAS and another set of 87 that were common between both cultivars TAS and SAS under aluminum stress (Fig. 2A). In the TAS cultivar, from the total miRNAs (191), 52% had been upregulated while in the SAS cultivar the majority of the miRNAs (75%) were down regulated (Fig. 2B). As can be seen in Fig. 2C, the cultivars had opposing expression profile. For the TAS cultivar the majority of the miRNAs (64%) were induced while in the SAS cultivar the majority (85%) were repressed (S1 Table). Generally, plant miRNAs can be classified into several different families where the members have similar sequences. The miRNAs identified in sugarcane roots belong to 100 known families (S1 Fig.) and among them, the most abundant miRNA families were miRNA159, miRNA156, miRNA 162, miRNA 396 and miRNA 444 (Fig. 3).

**Fig 2.**
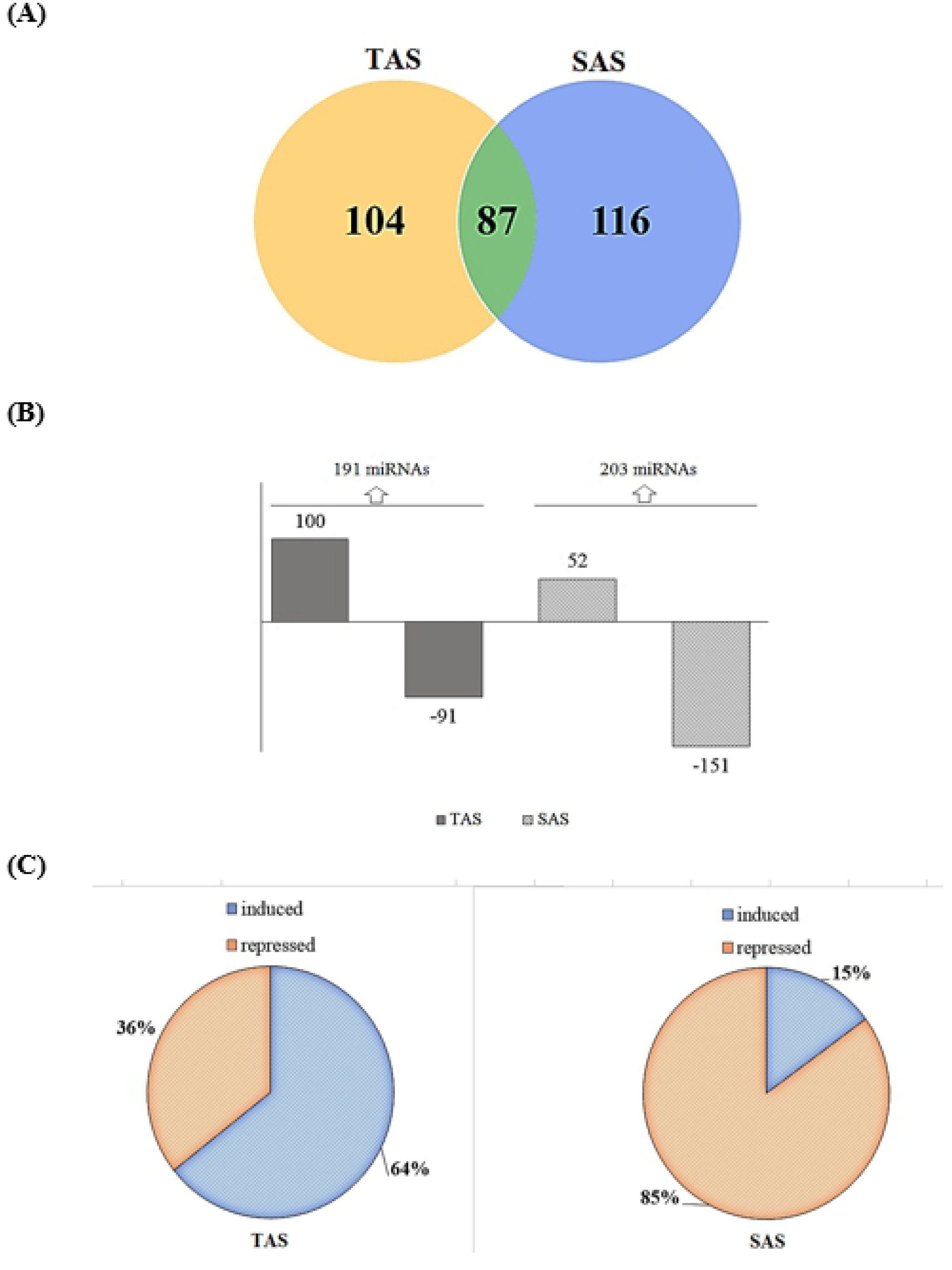
miRNAs expression profile. (A) Venn diagram showing miRNAs differently expressed in both cultivars; (B) The number of stress responsive miRNA is shown for each cultivar as well as the number of induced and repressed miRNAs under stress conditions; (C) Differential expression of the common miRNA between cultivars.

**Fig 3.**
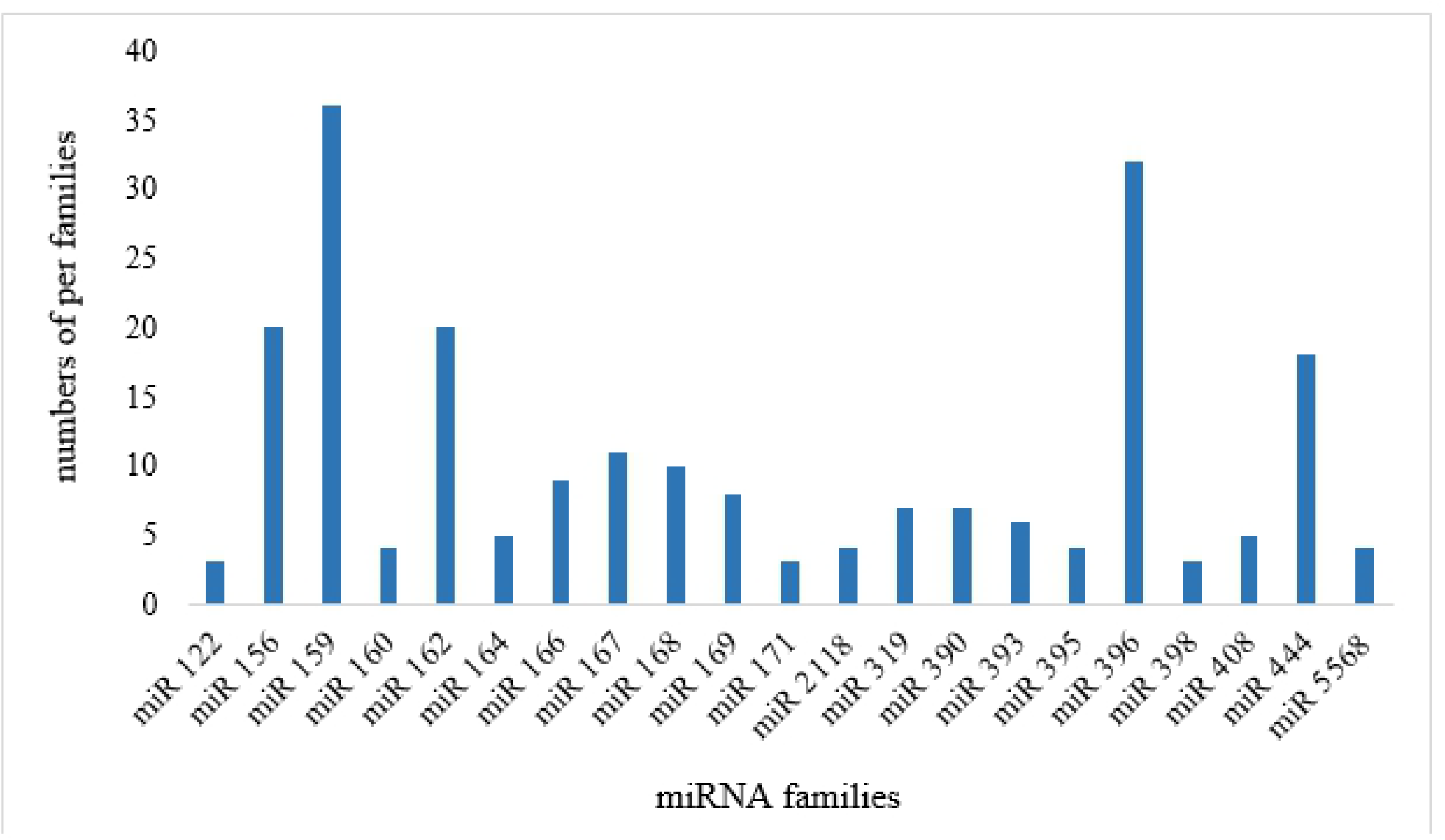
Most abundant miRNAs families identified in sugarcane roots.

Fourteen miRNAs were down-regulated, most of them in tolerant cultivar (TAS). Six were down-regulated in both cultivars (miR156, miR159, miR166, miR169, miR398, miR408) while three were down-regulated only in the sensitive cultivar. Two miRNAs showed to be up-regulated in the tolerant cultivar (miR168 and miR395) and contrasting expression was observed in 7 miRNAs (miR160, miR162, miR167, miR171, miR319, miR390, and miR396) Table 1.

**Table 1.** Expression analysis (Log2FC) of identified miRNAs in the sugarcane sequencing.

| miRNA Family |  | miRNA (reference) | Log2FC<sup>1</sup> |  |
| --- | --- | --- | --- | --- |
|  |  |  | TAS | SAS |
| Down-regulated | miR121 | miR121-1-npr (sit) | -4,88 | NR |
|  | miR122 | miR122-2-npr (sit) | -5,24 | NR |
|  | miR156 | miR156a-4 (sit) | -2,95 | -1,19 |
|  | miR159 | miR159a (sbi) | -2,95 | -1,01 |
|  | miR164 | miR164c (sit) | NR | -2,17 |
|  |  | miR164f-3p (zma) | -1,36 | NR |
|  | miR166 | miR166a-5p (zma) | -1,36 | -2,19 |
|  | miR169 | miR169n-5p (zma) | -1,36 | -1,19 |
|  | miR393 | miR393h (gma) | NR | -1,13 |
|  |  | miR393c-5p (zma) | -1,30 | NR |
|  | miR398 | miR398b-5p (zma) | -1,36 | -1,19 |
|  | miR408 | miR408 (csi) | -1,36 | NR |
|  | miR444 | miR444f (osa) | -2,36 | -1,59 |
|  | miR2128 | miR2128a-3p (gma) | -2,36 | NR |
|  | miR5568 | miR5568g-3p (sbi) | NR | -2,78 |
|  |  | miR5568f-3p (sbi) | -1,36 | NR |
|  | miR6253 | miR6253 (osa) | -2,30 | NR |
| Up-regulated | miR168 | miR168a-5p (zma) | 3,05 | NR |
|  | miR395 | miR395a (sly) | 4,85 | 1,12 |
| Constrasting | miR160 | miR160e-5p (osa) | -2,36 | 1,18 |
|  | miR162 | miR162b (ptc) | NR | 1,18 |
|  |  | miR162b (gma) | -1,36 | NR |
|  | miR167 | miR167h-3p (osa) | 4,29 | -4,36 |
|  | miR171 | miR171i (mdm) | 3,34 | -2,78 |
|  | miR319 | miR319-2 (sit) | 2,14 | -2,01 |
|  | miR390 | miR390a (cpa) | NR | 1,18 |
|  |  | miR390a (ath) | -1,36 | NR |
|  | miR396 | miR396d (zma) | 3,27 | -4,30 |

<sup>1</sup>NR: not responsive.

### miRNA transcriptome validation by RT-qPCR

RT-qPCR was used to validate the differentially expressed miRNAs through high-throughput sequencing in sugarcane roots. Six miRNAs (miR167, miR168, miR6253, miR159, miR156, miR121) modulated by aluminum were randomly selected for validation. The results of all these miRNAs confirmed by RT-qPCR were consistent with the high-throughput sequencing analyses (Fig. 4).

**Fig 4.**
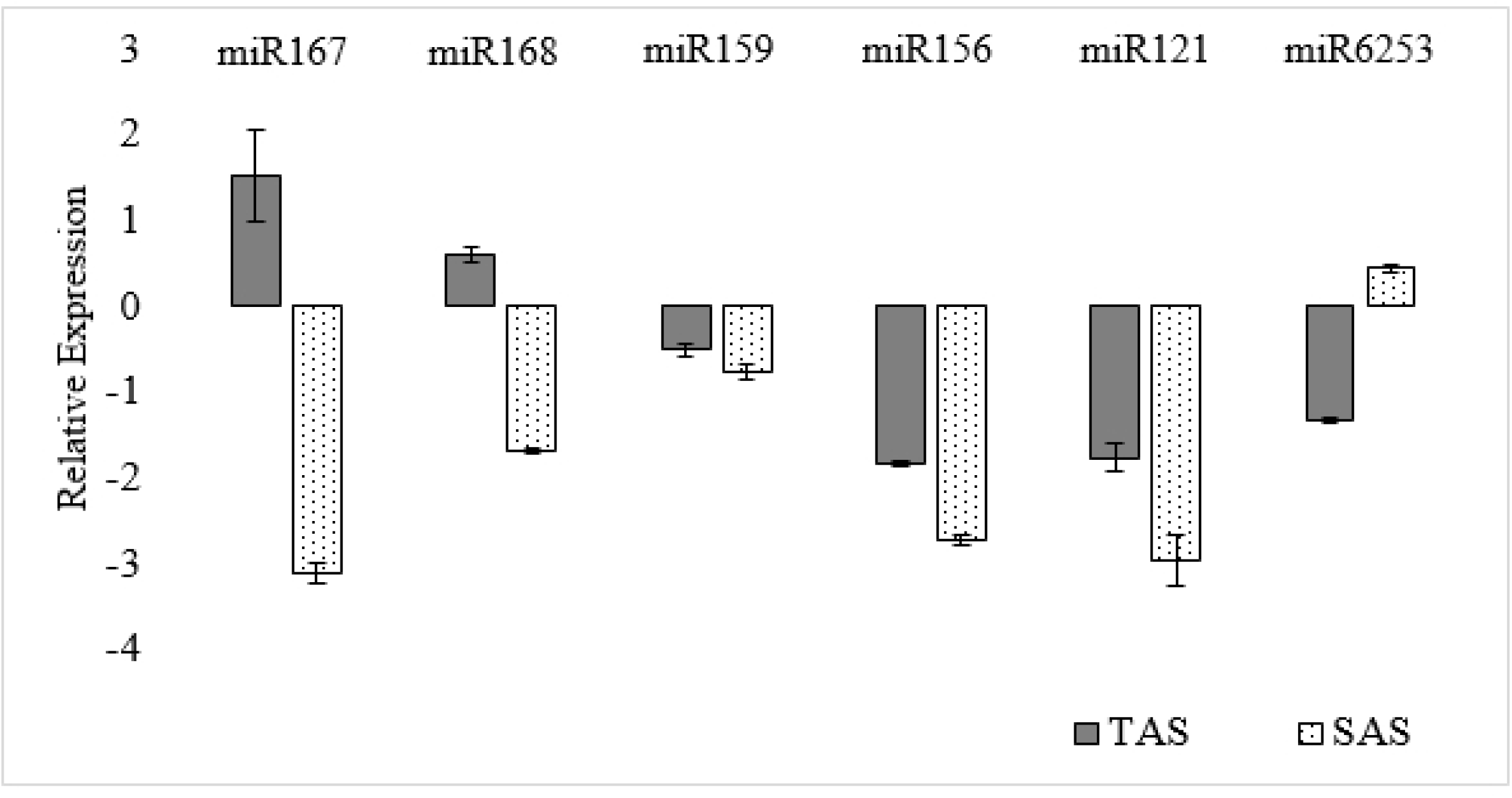
Relative expression of six identified miRNAs in sugarcane. Tolerant cultivar (TAS) and sensitive cultivar (SAS).

### Prediction of miRNA targets and GO annotation

Because plants miRNAs sequences are highly complementary to their targets, they can be used to predict their targets [37]. To better understand the possible biological function of the miRNAs the sequences of the most abundant microRNA families were used to search for their targets using Mercator, that assigns functional terms to nucleotide sequences (Table 2; S4 Table). The functional annotation of the targets is available in S4 Table. The genes and transcription factors regulated by the miRNAs participate in several biological processes: cell growth regulation (*LRR protein*), Auxin-activated signaling pathway (*Auxin response factor*), osmotic stress response (*CBL-interacting protein kinase 1*), growth negative regulation (*MYB domain protein 33*), among others.

**Table 2.** Predicted miRNA targets.

| miRNA | Potential targets by Mercator |
| --- | --- |
| 156 | *Squamosa promoter-binding protein-like* |
| 159 | *MYB domain protein; LRR protein* |
| 160 | *Auxin response factor* |
| 167 | *OsWAK; Copper-transporting ATPase PAA1* |
| 169 | *12-oxo-phytodienoic acid reductase* |
| 319 | *MYB domain protein* |
| 396 | *Growth-regulating factor* |
| 444 | *MADS-box transcription factor* |

## Discussion

Due to their regulatory role during plant development, the study of microRNAs associated to biotic and abiotic stress has increased significantly. Several miRNAs were identified in sugarcane in different tissues and stress conditions [29, 38] but none has been reported for sugarcane under aluminum stress. Here we report the first microtranscriptome associated with aluminum response in sugarcane. By comparing miRNA libraries sequences from the two contrasting cultivars, we were able to identify 394 differentially expressed miRNAs. Their size range from 17 to 25, with a majority between 20 and 24 nucleotides (Fig. 1), similar to the results reported for sugarcane under drought stress [29]. Small RNAs with different sizes may perform different functions. Twenty one nucleotide sRNAs are been associated to posttranscriptional gene silencing while 24nt mainly induce gene silencing by heterochromatin maintenance or RNA-Dependent DNA methylation [15, 39].

The contrast of the cultivars seen in the field was reflected in the microtranscriptome with opposing expression profile. For the tolerant cultivar (TAS) we observed that while 64% of microRNAs are been induced in the tolerant cultivar, in the sensitive the majority of microRNAs (85%) are been repressed under aluminum stress condition (Fig. 2C). Six of these miRNAs displayed the same expression profiles obtained by sequencing and RT-qPCR (Fig 4).

Those miRNAs were classified into different families (S1 Fig). The most abundant miRNA families were miRNA159, miRNA156, miRNA 162, miRNA 396 and miRNA 444 (Fig. 3). Members of those miRNA families has been identified in several crops associated to different stress conditions [27]. In our study, spp-miRNA156 was down-regulated in both cultivars (Table 3) it contains complementary sequences to *SQUAMOSA* (*SQUA*) *promoter*-*binding*-*like* (*SPL*) target gene which encode plant-specific transcription factors (Table 4). miR156 was induced in soybean, wheat [40] and repressed in rice [41] under drought, it was also identified in sugarcane under drought but it was not differently expressed. In Arabidopsis miR156 and its target *SPL3* were associated to the temporal regulation of shoot development [42]. In *Medicago truncatula* it was also down-regulated after 4 hours of aluminum stress [23] and it was classified as an early expressed gene.

miRNA159 was also down-regulated in both sugarcane cultivars by Al^+^ stress (Table 3). It targets a *MYB domain protein*, a transcriptional regulatory region (Table 4). miR159 has been associated with the control of multiple agronomic traits in rice [41], where it suppress cell division regulating negatively organ size. A transcriptome of a mutant suppressing miRNA159 revealed 7899 differentially expressed genes involved in several different pathways. Down-regulated genes were involved in pathways related to cell cycle, growth, signal transduction and hormone biosynthesis and signaling [41]. Although miR159 had been associated to aluminum stress in rice [41] and *Medicago* [23] it was not found in sugarcane before.

Three other miRNAs were down-regulated under aluminum stress: miR169, which targets a *12-oxo-phytodienoic acid redutase2*; miR398 a *Cooper/zinc superoxide dismutase*, involved in the cellular response to oxidative stress and miR444 that targets a *MAD-box transcription factor* associated to a wide range of functions including *e.g*. formation of flowers, flowering time control and vegetative development (Tables S3 and S4).

When we compared both cultivars TAS and SAS the miRNAs showed contrasting expression patterns under aluminum stress (Table 3). miRNA393h was down-regulated in the sensitive cultivar (SAS) and was not responsive in the tolerant cultivar (TAS) but miR393c, instead, showed to be down-regulated in the tolerant cultivar and not responsive in the sensitive. miR393 targets the *transport inhibitor response 1* gene (*TIR1*) (Table 3 and 4), required for normal response to auxin, essential for many important biological process in plants [43, 44], including root development [45]. One of the first symptom of Al^3+^ toxicity in plants is the reduction of lateral roots formation [46, 47]. Rice super expressing miRNA393a and miRNA393b shows a significant reduction in the lateral roots formation [48].

miRNA160 regulates the *Auxin response factor ARF* gene [49]. In our study, miRNA160 also showed a contrasting expression for the tested cultivars under Al^3+^ stress. It was down-regulated in the tolerant cultivar (TAS) and up-regulated in the sensitive (SAS) (Table 3). The repressed expression of miR160 in the TAS cultivar will increase the ARF (*Auxin response factor*) leading to the inhibition of lateral root formation. Increased concentrations of Al^3+^ also reduces the cytokine synthesis, transport, and increase abscisic acid concentration in roots [46, 50]. The same effect was observed in *Medicago truncatula* [23].

The predicted target for miR395 is the enzyme *Sulfate adenylyltransferase* (Table 4) important in adenosine 5’-phosphosulfate (ATPS) biosynthesis from ATP and inorganic sulfate [51]. It acts in the sulfate assimilation and reduction pathways in plants [52]. The up-regulation of miRNA395 in the TAS cultivar, can be an indicative that miR395 is modulating sulfur metabolic pathway as a response of increased Al^3+^ concentration. In acid soils sulfate absorption is increased [53], sulfate is normally reduced in the leaves but it can also be reduced in the roots [54] producing several compounds including glutathione playing important role in stress tolerance [55]. In sorghum, *ATPS1* and *ATPS2* genes were repressed under oxidative stress [52, 56]. In *Arabdopsis thaliana*, it was also demonstrated that miR395 is involved in the oxidative stress response modulating sulfur metabolic pathway [57].

Our results show that miRNA390 is down-regulated in TAS and up-regulated in SAS cultivar, it targets a *GTP-binding protein* (Tables 3 and 4). The repression of miR390 observed in TAS cultivar will lead to an increase in the expression of *GTP-binding protein.* Early signaling events in plant defense responses may involve ion channels, GTP-binding proteins and/or other signaling components [58]. It is well known that under adverse conditions, plant perceives a stress signal and transmits the information through signal transduction to the nucleus, resulting in altered physiological responses for surviving [59].

miRNA162 targets a *DICER-LIKE1 (DCL1)* an RNaseIII domain-containing protein responsible for the miRNAs synthesis [60, 61]. In our study miR162 was down-regulated in TAS and up-regulated in SAS. miRNA168 is also involved in the miRNA biogenesis targeting *Argonaute 1 AGO1* [62, 63]. miR162 and miR168 had been associated to the modulation of Cd stress in rice where both were down-regulated [63]. The authors suggested that the complexity of miRNA/target regulation and the altered expression of these miRNAs suggested that negative feedback regulatory circuits of the miRNA processing pathways might be highly active during Cd stress.

Our results show that in the TAS cultivar, the miRNAs 167, 171, 319 and 396 were up-regulated while in SAS they were down-regulated. miRNA171 and miR319 were also up-regulated in *M. truncatula* response to Al stress [21], the same for miRNA396 up-regulated in soybean in response to Al ([64]. In barley’roots, from XZ29 a genotype Al-tolerant, under aluminum stress, miRNA 319 was up-regulated while miRNA396 was down-regulated [65].

miRNAs 171 and miR396 have been reported as part of the answer to abiotic stress regulation [66]. Under Al^3+^ stress, some genes such as *ARF*, domain-containing *Cation-transporting ATPase* and *MYB*, were found to be cleaved in soybean [64]. The target genes for miRNAs 167 e 319 are *ATPase activity* and *MYB domain*, respectively, act on ion homeostasis, negative regulation of growth and positive regulation of abscisic acid-activated signaling pathway. Aluminum can trigger protective mechanisms involving miRNAs that can improve the plant’s tolerance to Al toxicity [64].

## Conclusions

This is the first study of global identification of miRNAs responsive to aluminum stress in contrasting sugarcane cultivars. The study provides a basis for the understanding of molecular mechanisms associated with tolerance in sugarcane under aluminum stress indicating miRNAs that modulate specific target genes involved in roots development and plant aluminum stress response.

## Supporting information

**S1 Fig. Summary of identified miRNA families in sugarcane and number of miRNAs per family.**

(TIF)

**S1 Table. Stress-responsive miRNAs identified in TAS and SAS.**

(DOCX)

**S2 Table. The primer sequences used in the qRT-PCR validation.**

(DOCX)

**S3 Table. miRNAs sequences evaluated.**

(DOCX)

**S4 Table. Distribution of predicted miRNA targets genes**. Functional annotation of target genes regulated by the most abundant miRNA families differentially expressed.

(DOCX)

## AuthorContributions

**Conceived and designed the experiments:** SMZ; KRK;

**Performed the experiments:** RGS; TMR;

**Analyzed the data:** SMZ; RGS; TMR; KRK; PK;

**Contributed reagents/materials/analysis tools:** SMZ; KRK;

**Wrote the paper:** RGS; SCF; SMZ;

